# On the Use of Double Mutant Cycles to Probe the Molecular Interactions in Biomolecular Condensates

**DOI:** 10.64898/2026.02.03.703500

**Authors:** Arriën Symon Rauh, Giulio Tesei, Kresten Lindorff-Larsen

## Abstract

Disordered proteins can form biomolecular condensates by demixing from their environment, enabling reversible compartmentalisation of cellular components in the form of membraneless organelles. Multivalent interactions are essential for this type of phase separation behaviour, and for disordered proteins, the potential for multivalent interactions is encoded in the sequence composition and patterning. Mutational studies have been instrumental in helping elucidate this sequence grammar by perturbing the amino acid sequence and quantifying the resulting changes in the driving force for phase separation. While such studies have provided a detailed and predictive understanding of the driving forces for phase separation, they strictly do not inform on the nature of the interactions that drive phase separation. Here, we propose using double mutant cycles to explore molecular interactions and their contributions to condensate properties more directly. We explore the applicability of double mutant cycles for different types of interactions in condensates formed by the low-complexity domain of hnRNPA1 using coarse-grained molecular dynamics simulations. We find that the interactions between arginine and tyrosine residues, as well as between aromatic residues, contribute mostly additively to the propensity for phase separation. However, for the interactions between charged residues, we find that—in an interplay with the net charge of the protein—there is a measurable non-additive contribution to the phase separation propensity. Based on our results, we envisage that double mutant cycles could provide additional insights into protein phase separation, thus expanding the understanding of the sequence grammar and the underlying molecular interactions.

## Introduction

Both intrinsically disordered proteins and intrinsically disordered regions (here, collectively called IDRs) may exhibit phase separation (PS) behaviour that can lead to the formation of biomolecular condensates or membraneless organelles (***Brangwynne et al., 2015; Banani et al., 2017; Holehouse and Kragelund, 2023; Pappu et al., 2023***). This process involves the demixing of protein chains from their environment, forming concentrated phases of either one type of protein (homotypic PS) or forming concentrated phases that are more complex mixtures (heterotypic PS) of for example multiple different proteins and RNA. In this context, IDRs can function both as a driver for PS or as a client that gets recruited into a condensate.

The formation of biomolecular condensates may be modulated by a variety of physico-chemical and environmental factors such as temperature, pH, salt concentrations and macromolecular crowding. However, experiments, theory and simulations have demonstrated that the intrinsic propensity of an IDR to exhibit PS behaviour is fundamentally tied to its solubility and ability to form multivalent interactions and percolated interaction networks (***Pappu et al., 2023***). For disordered proteins, this potential for multivalent interactions is encoded in the sequence composition and the patterning of the different types of amino acid residues (e.g., charged or aromatic) (***Lin et al., 2018; Dignon et al., 2018b; Martin and Mittag, 2018; Martin and Holehouse, 2020; Borcherds et al., 2021***). To describe the sequence composition and patterning, a variety of measures have been conceived (***Das and Pappu, 2013; Sawle and Ghosh, 2015; Zheng et al., 2020a; Ghosh et al., 2022; Ruff et al., 2025***). For instance, *κ* (***Das and Pappu, 2013***) and the sequence charge descriptor (SCD) (***Sawle and Ghosh, 2015***) describe the patterning of charged residues, and the sequence hydropathy descriptor (SHD) (***Zheng et al., 2020a***) informs on the patterning of hydrophobic residues along the sequence. Other parameters quantify aspects of the sequence composition, such as the fraction of charged (FCR) or aromatic (*f*_aro_) residues and the net charge per residue (NCPR).

Experiments have—often in concert with theory and simulations—been used to help elucidate the sequence properties that help drive homotypic PS of IDRs. For example, an extensive study of the PS behaviour of a group of prion-like RNA binding proteins such as Fused in Sarcoma (FUS), (***Wang et al., 2018***) showed that tyrosine residues in the prion-like domain (PLD) and arginine residues in the RNA binding domain (RBD) are important, and that the PS behaviour can be rationalised quantitatively by a theory of associative polymers (***Rubinstein and Dobrynin, 1997; Semenov and Rubinstein, 1998***) using arginine and tyrosine residues as ‘stickers’ (***Wang et al., 2018***). For the low-complexity domain of hnRNPA1 (A1-LCD), mutational experiments revealed that the number of aromatic residues, to a large extent, dictates the PS propensity, whereas the patterning of aromatic residues influences how aggregation-prone the sequence is (***Martin et al., 2020***). Additional experiments and analyses of A1-LCD revealed additional effects of other residue types, including the importance of NCPR for modulating PS (***Bremer et al., 2022***). Experiments based on an mRNA display method have recently been used to study and quantify the driving force for hetero- and homotypic PS across thousands of IDRs, thus helping to consolidate these rules across a wider range of sequence space (***Norrild et al., 2025***).

These types of experiments on FUS, A1-LCD, and several other systems have, together with computational methods, resulted in a relatively good understanding of the general rules that determine the extent to which an IDR will undergo PS in vitro (***Pappu et al., 2023***). The outcomes of mutational studies have thus been used to suggest the importance of different types of molecular interactions that contribute to condensate formation such as the hybrid cation–*π*/*π*–*π* interactions between arginine and tyrosine residues (***Nott et al., 2015; Pak et al., 2016; Boeynaems et al., 2017; Wang et al., 2018; Bremer et al., 2022***). However, even with a good understanding of the residue types and sequence properties that determine PS, it is difficult to determine the molecular nature of the interactions that underlie these rules. This is in part because the interplay and balance between protein–protein and protein–solvent interactions are complex and system dependent, making it difficult to link the sequence properties with specific types of interactions mediating IDR PS. Nevertheless, some of the types of interactions that have been suggested to be important include cation-*π* interactions between cationic arginine and lysine residues and aromatic residues like phenylalanine, tyrosine and tryptophan (***Dougherty, 1996; Gallivan and Dougherty, 1999; Dougherty, 2013; Mahadevi and Sastry, 2013***). In this case, arginine has a special character relative to lysine where the guanidinium group acts as a *π*-system allowing for *π*–*π* interactions with aromatic residues (***Burley and Petsko, 1988; Gobbi and Frenking, 1993***). Besides this, *π*–*π* interactions formed, for example between aromatic residues, have been suggested to be important for the formation of biomolecular condensates (***Vernon et al., 2018***). Electrostatic interactions have also been shown to be potential drivers of PS and allow for encoding salt-dependent effects (***Zhou and Pang, 2018; Boyko et al., 2019; Galvanetto et al., 2023***). Finally, hydrogen bonds formed by the protein backbone or side chains have been implicated in modulating PS (***Murthy et al., 2019; Guo et al., 2022***). For example, hydrogen bonds between low-complexity aromatic-rich kinked segments (LARKS) have been suggested to be associated with transiently formed beta-sheet-rich structures in biomolecular condensates, which in turn have been associated with condensate ‘ageing’ and fibril formation processes (***Hughes et al., 2018, 2021; Blazquez et al., 2023***).

To study the molecular interactions underlying the PS behaviour of proteins, experimental methods such as NMR spectroscopy, single-molecule Fluorescence Resonance Energy Transfer (smFRET), fluorescence microscopy in combination with microfluidic devices, and other spectroscopic approaches have been applied (***Arosio et al., 2016; Murthy et al., 2019; Martin et al., 2020; Murthy and Fawzi, 2020; Ganser and Myong, 2020; Arter et al., 2020; Linsenmeier et al., 2021; Beck et al., 2024***). NMR spectroscopy can, for example, be used to obtain insights into interacting residues of IDRs both outside of and inside condensates (***Murthy et al., 2019; Martin et al., 2020; Kim et al., 2021; Emmanouilidis et al., 2021; Guseva et al., 2023***), and smFRET can be used to probe which regions of IDRs make intramolecular interactions (***Joshi et al., 2023; Galvanetto et al., 2023***). Finally, cryogenic-sample electron tomography has the potential to provide information on interaction networks in condensates through a three-dimensional structure (***Rizvi et al., 2021; Scholl et al., 2024; Zhou et al., 2025***).

Molecular simulations and theory complement experiments and have been used extensively to study the interactions and dynamics within condensates. A range of different approaches have been used to tackle the complexity, size and slow dynamics of condensates; these range from allatom molecular dynamics (MD) simulations (***Ivanović and Best, 2025***) to coarse-grained models (***Maristany et al., 2025; Rizuan et al., 2026***). All-atom simulations have, for example, been used to quantify contacts within condensates (***Rekhi et al., 2024***), the location of ions (***Zheng et al., 2020b***), and show rapid exchange between competing ionic contacts (***Galvanetto et al., 2023***).

Residue-level coarse-grained (CG) MD simulations have also been used to guide and interpret PS experiments. By encoding hypotheses about the importance of different types of interactions and their relative strengths, these models can be used to examine the physico-chemical basis for PS (***Dignon et al., 2020***). For example, ***Das et al***. (***2020***) evaluated the effects of adding specific terms for cation–*π* interactions to the HPS model by comparing to experimental PS data. We and others have taken a top-down approach to parameterise IDR models against experimental data (***Norgaard et al., 2008; Martin et al., 2020; Latham and Zhang, 2020; Dannenhoffer-Lafage and Best, 2021; Tesei et al., 2021***) and thereby to provide a data-driven set of interaction parameters that quantify the strengths of the molecular interactions. Given the coarse-grained nature of these models as well as the limited amounts of data, it is nevertheless challenging to use these to define the importance of individual types of interactions uniquely. For example, some models represent amino-acid interactions with specific terms for different pairs of amino acid types (***Latham and Zhang, 2020; Joseph et al., 2021; Bremer et al., 2022***), whereas other models only have one parameter per amino-acid type, which are then combined to describe the pairwise interactions (***Dignon et al., 2018b; Tesei et al., 2021***). Similarly, we have shown that a simple Debye-Hückel model for electrostatics can be used to capture some aspects of the salt-dependence of PS of A1-LCD (***Tesei and Lindorff-Larsen, 2023***) and how PS depends on NCPR (***Tesei et al., 2021***), but such a model does not capture all of the complex electrostatic effects that occur in condensates (***Posey et al., 2024; Dai et al., 2024***).

Double mutant cycles provide a potent mutational approach for the more direct quantification of the contribution of individual interactions to an overall measurable property, such as folding stability or catalytic activity (***Jencks, 1981; Carter et al., 1984; Fersht et al., 1992; Schreiber and Fersht, 1995; Horovitz, 1996; Horovitz et al., 2019***). A double mutant cycle entails identifying a set of interacting amino acid residues and measuring, for example, the thermodynamic stability for the wild-type (WT) protein, two single mutants, and a double mutant where the two single mutants are combined (Fig. 1A). The difference between the sum of the two individual perturbations and the effect of the double mutant can—under certain assumptions—provide an estimate of the direct interaction between two amino acid residues (***Fersht et al., 1992***). This difference is often termed a coupling or interaction energy (ΔΔ*G*_int._), and when there is no coupling, the interaction can be interpreted in an additive manner (***Wells, 1990; Skinner and Terwilliger, 1996***). This approach has been applied to study the contributions of aromatic–aromatic interactions (***Serrano et al., 1991***), ionic (***Serrano et al., 1990; Šali et al., 1991***) and hydrogen bonds (***Marqusee and Sauer, 1994***) to protein stability, and has recently been extended to larger scales using multiplexing (***Araya et al., 2012; Olson et al., 2014; Diss and Lehner, 2018; Tsuboyama et al., 2023***).

**Figure 1.**
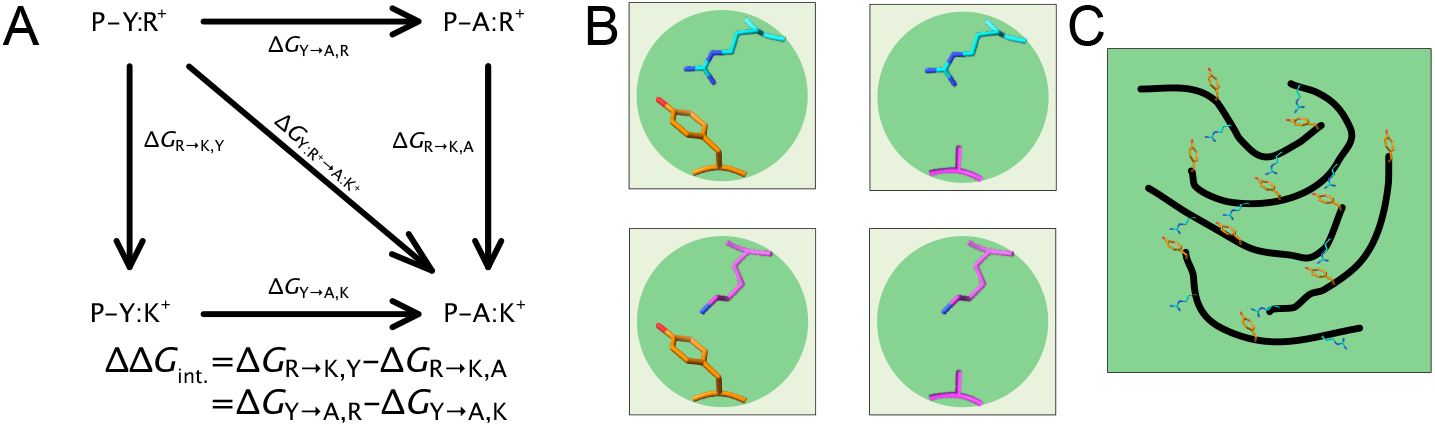
Outline of the double mutant cycle approach and the application to phase separation. (A) An example of a double mutant cycle for tyrosine-arginine interaction is when the tyrosine residues are substituted with alanines and arginines with lysines. (B) Molecular interactions for the wild-type interaction, the single mutants and the double mutant. (C) Illustration of the complexity of the interactions in the case of homotypic PS, where multiple chains of an IDR phase separate together, resulting in multiple cases of the same interaction, e.g. tyrosine–arginine.

While the interpretation of double mutant cycles is easiest when only two single amino acid changes are combined and when the perturbed interactions are structurally well-defined (Fig. 1B), the formalism can be applied more broadly. Previous mutational experiments of PS have reported results in which two different sets of residues are studied individually and combined (***Wang et al., 2018; Garcia-Cabau et al., 2025***). For example, ***Wang et al. (2018***) combined Tyr-to-Phe mutations in the FUS PLD with Arg-to-Lys mutations in the FUS RNA binding domain, though it did not directly quantify the extent to which these perturbations were additive.

The goal of our work is to explore whether double mutant cycles could be used to quantify the contribution of particular types of molecular interactions in biomolecular condensates. To do so, we have used our coarse-grained CALVADOS 2 model (***Tesei and Lindorff-Larsen, 2023; von Bülow et al., 2025***) to explore what outcomes could be expected. We probe three types of interactions; the hybrid cation–*π*/*π*–*π* interactions between Arg and Tyr residues that have been suggested to be of importance for the PS of FUS-like IDRs (***Nott et al., 2015; Pak et al., 2016; Wang et al., 2018***), the *π*– *π*-interactions between aromatic sticker residues that have been shown to be the most interacting residues in A1-LCD condensates (***Martin et al., 2020***) and charge–charge interactions in A1-LCD (***Bremer et al., 2022***). In the cases we examined, we mostly find couplings between the interactions, suggesting they contribute additively. We observe a significant coupling only for the case of charge– charge interaction, where the overall net charge of the sequence is modified.

## Results and Discussion

### Interrogating tyrosine–arginine interactions in biomolecular condensates

Our understanding of the driving forces for PS of IDRs is, to a large extent, based on using mutations to perturb the sequence and probe the effect on PS. In this manner, for example, the sotermed ‘sticker’ residues can be ranked based on the effect a mutation has on the saturation concentration (*c*_sat_) or a partitioning free energy (Δ*G*_trans_ = *RT* ln *c*_dil_/*c*_den_, where *c*_dil_ = *c*_sat_ when there is PS). Ranking or determining the importance of the specific interactions found in a biomolecular condensate is less trivial. For structured proteins, double mutant cycles have been used to probe the contribution of specific interactions to, for example, thermodynamic or kinetic properties by exploring if the effect of mutations is non-additive (***Schreiber and Fersht, 1995; Horovitz, 1996***).

We began by exploring the contribution of the interaction of the sticker residues tyrosine and arginine, which has been suggested to be one of the stabilising interactions for IDR-mediated condensates of PLDs, given the hybrid cation–*π*/*π*–*π* character (***Wang et al., 2018; Farag et al., 2022; Rekhi et al., 2024***). We chose to use the well-studied homotypic PS of A1-LCD as the CALVADOS model has been shown to perform well in predicting PS properties of the WT and a variety of variants (***Tesei and Lindorff-Larsen, 2023; Pesce et al., 2024; Von Bülow et al., 2025***). We manually designed single and double mutants perturbing potential Tyr–Arg interactions by selecting five residues of each type that are found to make Tyr–Arg interactions in the condensates (Fig. S1A). We note here that we use the nomenclature ‘single mutant’ even when multiple residues are substituted simultaneously as a single group; similarly, we use the term ‘double mutant’ when two such single mutants are combined. We chose to substitute both tyrosine and arginine residues in multiple locations in the single mutants to obtain a substantial and measurable change in PS (Fig. 2A and Table S1). Specifically, we chose to substitute arginine to lysine residues (Δ*λ* = –0.55) to maintain the net charge of the protein, and to substitute the strong tyrosine sticker residues with the slightly less sticky phenylalanine residues (Δ*λ* = –0.11, where *λ* refers to the CALVADOS stickiness scale) so as not to destabilise the double mutant too much.

**Figure 2.**
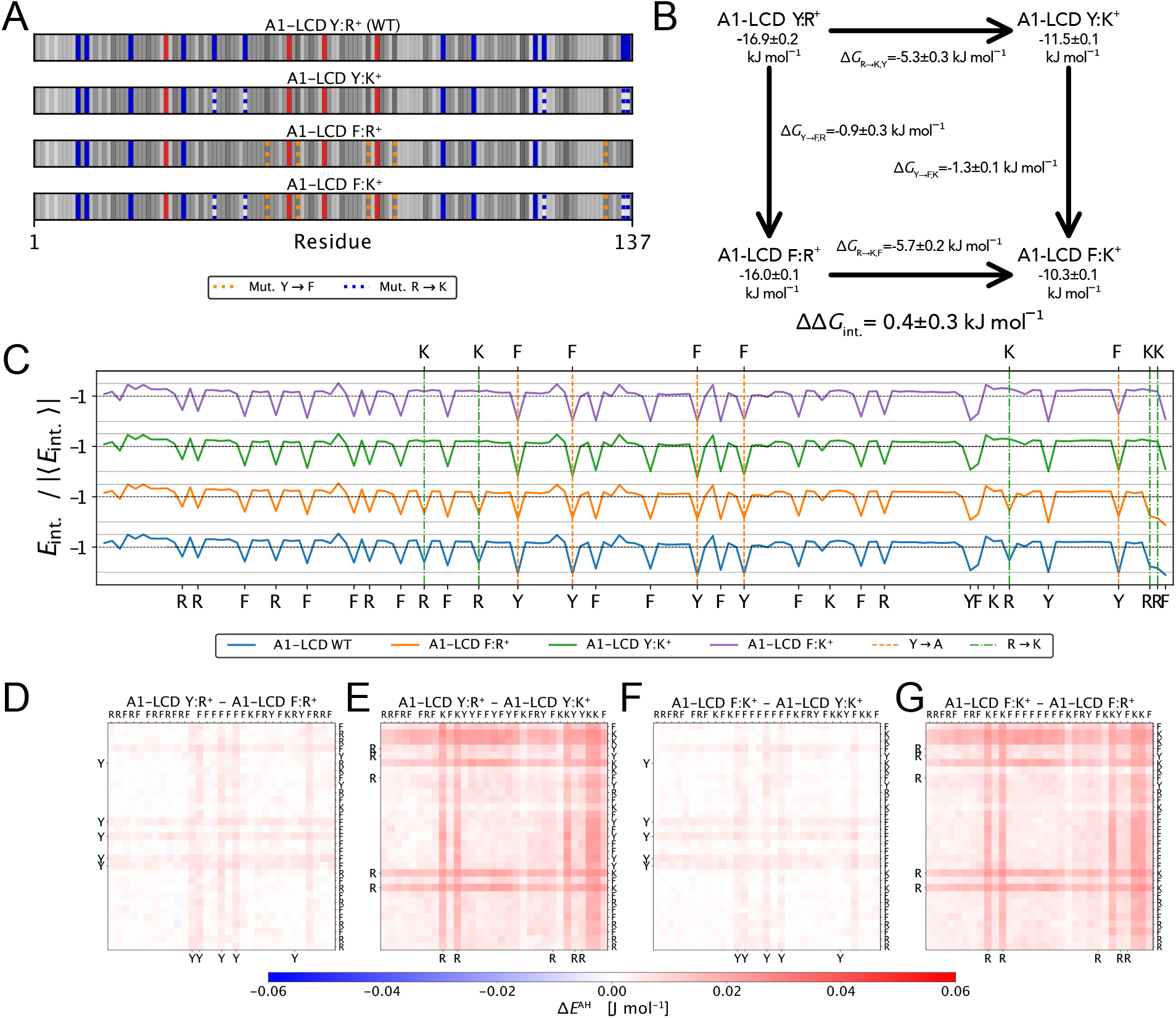
Probing the interactions between tyrosine and arginine residues with a double mutant cycle. (A) Sequence plots of WT A1-LCD and variants. The blue and red lines represent positively and negatively charged amino acid residues, respectively. The neutral amino acids are coloured in greyscale to represent their hydropathy based on the CALVADOS 2 *λ* parameters. The dashed lines represent the locations where we substituted tyrosine with phenylalanine residues (orange) and arginine with lysine residues (blue). (B) Double mutant cycle for perturbing Arg–Tyr interactions with the mutations R→K and Y→F. Free energies of phase separation are indicated for each variant, and the changes in free energies are shown next to the arrows. (C) From the bottom to top, 1D projections of the energy maps for (blue) WT A1-LCD, (orange) Y→F, (green) R→K, and (purple) Y→F & R→K. The maps are normalised by the absolute average interaction energy *E*. Difference maps for sticker residue interaction energies of (D) substituting tyrosines, (E) substituting arginines, (F) substituting tyrosines after substituting arginines, and (G) substituting arginines after substituting tyrosines.

We performed direct co-existence simulations in a ‘slab’ geometry of the A1-LCD WT, A1-LCD Y:K^+^ (5 R→K), A1-LCD F:R^+^ (5 Y→F), and the double mutant A1-LCD F:K^+^. Our simulations show that both of the single mutants (Δ*G*_trans_ –11.5 ± 0.1 kJ mol^−1^ & –16.0 ± 0.1 kJ mol^−1^) and the double mutant (Δ*G*_trans_ –10.3 ± 0.1 kJ mol^−1^) are less prone to phase separate than WT (Δ*G*_trans_ –16.9 ± 0.2 kJ mol^−1^) (Fig. 2B). From these PS propensities, we calculate the change in propensity for the single mutations (Δ*G*_R→K,Y_ and Δ*G*_Y→F,R_) and for the change in propensity when performing the same mutations after an initial mutation (Δ*G*_R→K,F_ and Δ*G*_Y→F,K_). From these we calculate the coupling energy for the Tyr– Arg interactions as the difference between the two vertical (or equivalently the two horizontal) legs of the thermodynamic cycle (eg. ΔΔ*G*_int_ = Δ*G*_Y→F,R_ – Δ*G*_Y→F,K_). The calculated coupling energy is relatively small (ΔΔ*G*_int._ = 0.4 ± 0.3 kJ mol^−1^), suggesting that there is no sizeable coupling between the two interacting sets of residues, which suggests that the Tyr–Arg interactions are additive in their contribution to the stabilisation of the biomolecular condensate of A1-LCD.

We then explored the changes in molecular interactions when perturbing the Tyr–Arg interaction by analysing the non-ionic residue–residue interaction maps of the WT and variants (Fig. S1). From the interaction maps we observe that, as described previously (***Martin et al., 2020; Tesei et al., 2021; Bremer et al., 2022***), the sticker residues tyrosine, arginine and phenylalanine have the most favourable contributions (lowest 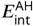) and the most significant changes in interactions occur at the positions where we substitute these residues (Fig. 2C). The 1D projections of the energy maps suggest that there may be some compensatory increase in interactions in alternative sticker positions. To investigate this further, we calculated the interaction energy difference maps for the four different mutational steps, focusing on the tyrosine, arginine, phenylalanine and lysine residue positions (Fig. 2D–G). Overall, these difference maps are consistent with the Δ*G*-values calculated for the double mutant cycle. The maps representing the tyrosine to phenylalanine substitutions (Fig. 2D & F) and those representing the arginine to lysine substitutions (Fig. 2E & G) are similar, and the energy difference is higher in the latter case. We do not observe any clear compensatory formation of new interactions in non-mutated positions along the sequence, though we note that the need for normalisation of the maps makes comparisons difficult.

Because both tyrosine and phenylalanine are strong stickers, the Tyr-to-Phe substitutions have a relatively modest effect on PS (Fig. 2B), potentially masking any coupling energy present. As an alternative, we thus also performed a double mutant cycle by changing the tyrosine to alanine residues at the same positions (Fig. S2A). As the Y→A substitution comes with a much larger difference in *λ*, the double mutant did not form a stable condensed phase in the simulations and thus prevented us from determining Δ*G* directly. To estimate the PS propensity of the double mutant, we instead used a computational polyethylene glycol (PEG) titration (***Rauh et al., 2025***) to estimate an extrapolated value of 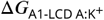 without PEG (Fig. S2B). When calculating the coupling energy from this modified double mutant cycle, we observe a similarly low value (ΔΔ*G*_int._ = 0.1 ± 0.4 kJ mol^−1^), suggesting no coupling and no specific contribution of the Y–R interactions (Fig. S2C).

### Perturbing the number and patterning of aromatic residues

After establishing that the potential *π*-*π*/cation-*π* interactions between tyrosine and arginine residues do not exhibit a strong coupling in our simulations, we set out to explore whether we could detect a coupling in *π*-*π* interactions between aromatic residues along the sequence of A1-LCD. In particular, it has been shown in experiments and simulations that both the number of aromatic residues and how they are distributed along the sequence dictate the PS behaviour of A1-LCD (***Martin et al., 2020***).

To probe for coupling between the aromatic residues with a double mutant cycle, we designed two sets of A1-LCD variants perturbing the aromatic residues at the termini and in the central ‘core’ region of the sequence (Fig. 3A and Table S1). We made these two sets of variants to examine whether there may be differences due to residue proximity along the sequence. Specifically, we have previously shown that residues near the termini form more interactions in condensates compared to in the dilute phase (***Tesei et al., 2021***), so we speculated that they might behave differently in the mutational studies.

**Figure 3.**
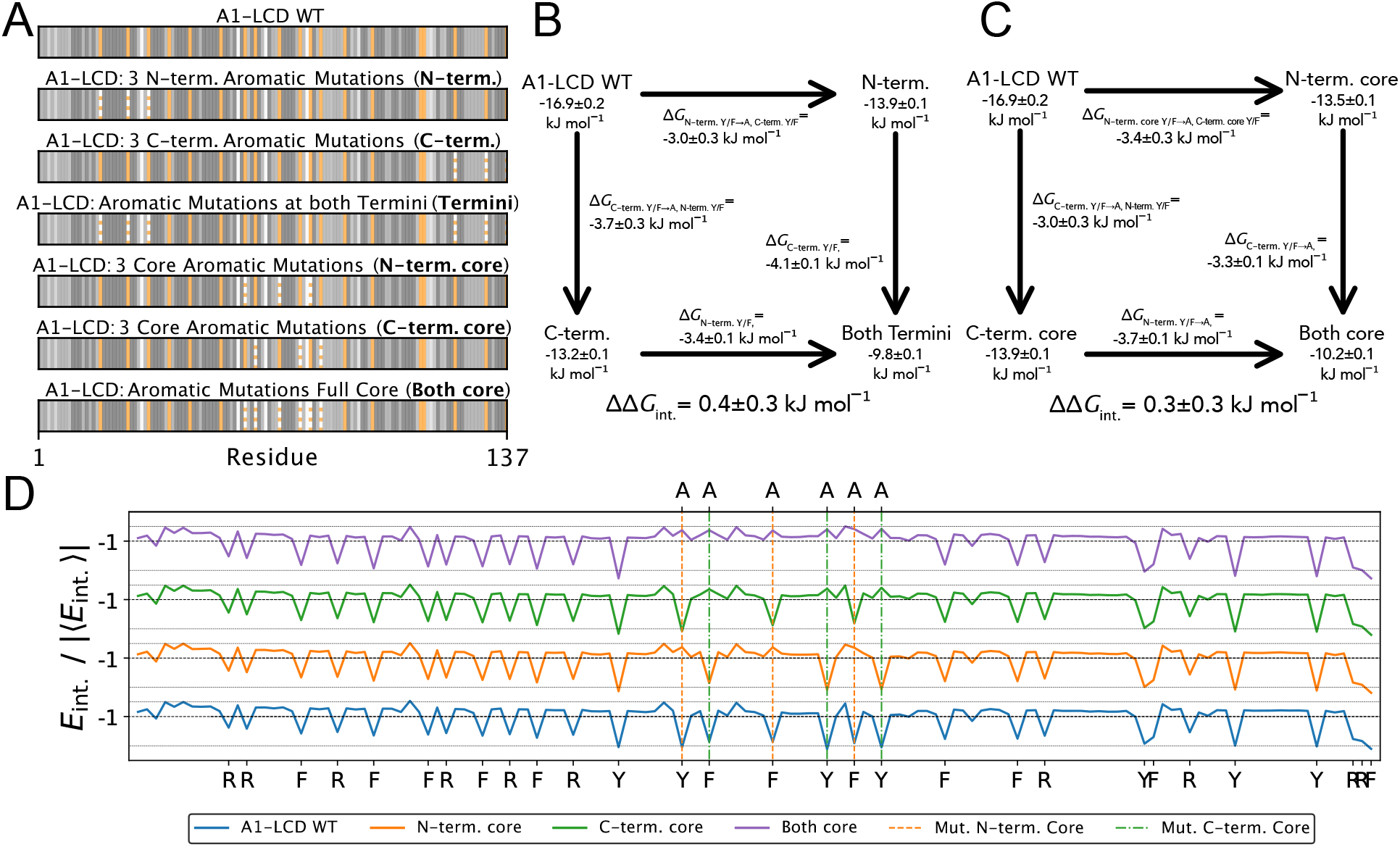
Probing interactions between aromatic residues in different sequence regions. (A) Sequence plots of A1-LCD variants perturbing the number and patterning of aromatic residues. The neutral amino acids are coloured in greyscale to represent their hydropathy based on the CALVADOS 2 *λ* parameters. The solid orange lines represent the tyrosine and phenylalanine residue positions, and the dashed orange lines are where any of those are substituted with alanine residues. (B) Double mutant cycle for perturbing aromatic residues at the termini with substitutions to alanine residues. (C) Double mutant cycle for perturbing aromatic residues in the central ‘core’ region of the sequence with substitutions to alanine residues. (D) From the bottom to top, 1D projections of the energy maps for (blue) A1-LCD WT, (orange & green) A1-LCD where two different sets of aromatic residues in the core are mutated, and (purple) A1-LCD where all core aromatics are mutated. The original maps are normalised by the absolute average interaction energy |⟨*E*⟩ |.

In the first set, we substituted three aromatic residues at the N-terminal part and three at the C-terminal part of the sequence. In the second set, we substituted six aromatic residues in the central region of the sequence. With all these variants, we only modify the overall hydropathy of the sequence ( *λ*) and the patterning of aromatic residues, without changing the net charge or the charge patterning (Table S2). We then used slab simulations of the variants to determine the energetic values for the double mutant cycles. The results show that the single mutants from both groups of variants show a similar propensity to phase separate, with in all cases the same number of aromatic residues substituted (Fig. 3B and C). The small difference in Δ*G*_trans_ between the single mutants in the different groups can be explained by the numbers of tyrosine versus phenylalanine residues being substituted (2:1 for the ‘N-term.’ and ‘N-term. core’ single mutants and 1:2 for the ‘C-term.’ and ‘C-term. core’ single mutants). This difference is then also visible in the mutational Δ*G*-values, where mutating more tyrosine residues results in a greater change. However, we find no sizeable coupling (ΔΔ*G*_int._ = 0.4 ± 0.3 kJ mol^−1^ and 0.3 ± 0.3 kJ mol^−1^; Fig. 3B and C). Thus, again, we find no evidence from the estimated couplings for any specific interactions, even though the simulations clearly show strong interactions between the mutated sites in the WT sequence.

We visualise the impact of the perturbations at the molecular interaction level by analysing the residue–residue interaction maps (Fig. S3A–D and Fig. S3A–D). The 1D projections of the interaction maps show that the aromatic and arginine residues form the most stable interactions, and that at the positions where we introduced substitutions, the interaction energy drops to a baseline (Fig. 3D and Fig. S4I). The interactions appear to become more stable in sticker positions that are not mutated, but the overall baseline also shifts. To explore this further, we calculated the difference maps, focusing only on positions with sticker residues. These maps also demonstrate that the differences in interaction energies are overall similar between the single mutant and double mutant steps for both sets of variants, in line with the mutational Δ*G*-values based on the PS propensities (Fig. S3E-G and Fig. S4E-G). While in the case of the double mutant, the already mutated positions show less destabilisation, overall, all the sticker–sticker interactions are destabilised, and there appears to be no substantial compensatory increases.

### Perturbing the net charge and charge patterning of a designed A1-LCD variant

We next used double mutant cycles to address the contribution of charge–charge interactions. To this end, we chose to use as a starting point a recently described redesigned variant of A1-LCD (termed ‘V1’). In particular, V1 has the same amino acid composition as A1-LCD, but is more compact and has a stronger driving force for PS as a result of the segregation of oppositely charged residues to the termini (Fig. 4A) (***Pesce et al., 2024***). These charge groups form strong interactions in simulations of V1 condensates (***Pesce et al., 2024***), and so we hypothesised that we might be able to detect these interactions using a double mutant cycle. We designed two groups of variants perturbing different aspects of the charge interactions. In the first set, we substituted both positive residues (five arginines and one lysine) and negative residues (four aspartates) with alanine residues (Fig. 4A and Table S1). In this manner, we perturb both the charge patterning and the capability to form intra- and inter-molecular charge–charge interactions; however, we also change the net charge of the protein so that each of the four variants has a different net charge. As an alternative approach to perturb the long-range charge interactions without modifying the net charge, we swap the positions of the selected charged residues with glycine, serine or alanine residues positioned along the sequence, but not at the termini, thus retaining the amino acid composition and net charge.

**Figure 4.**
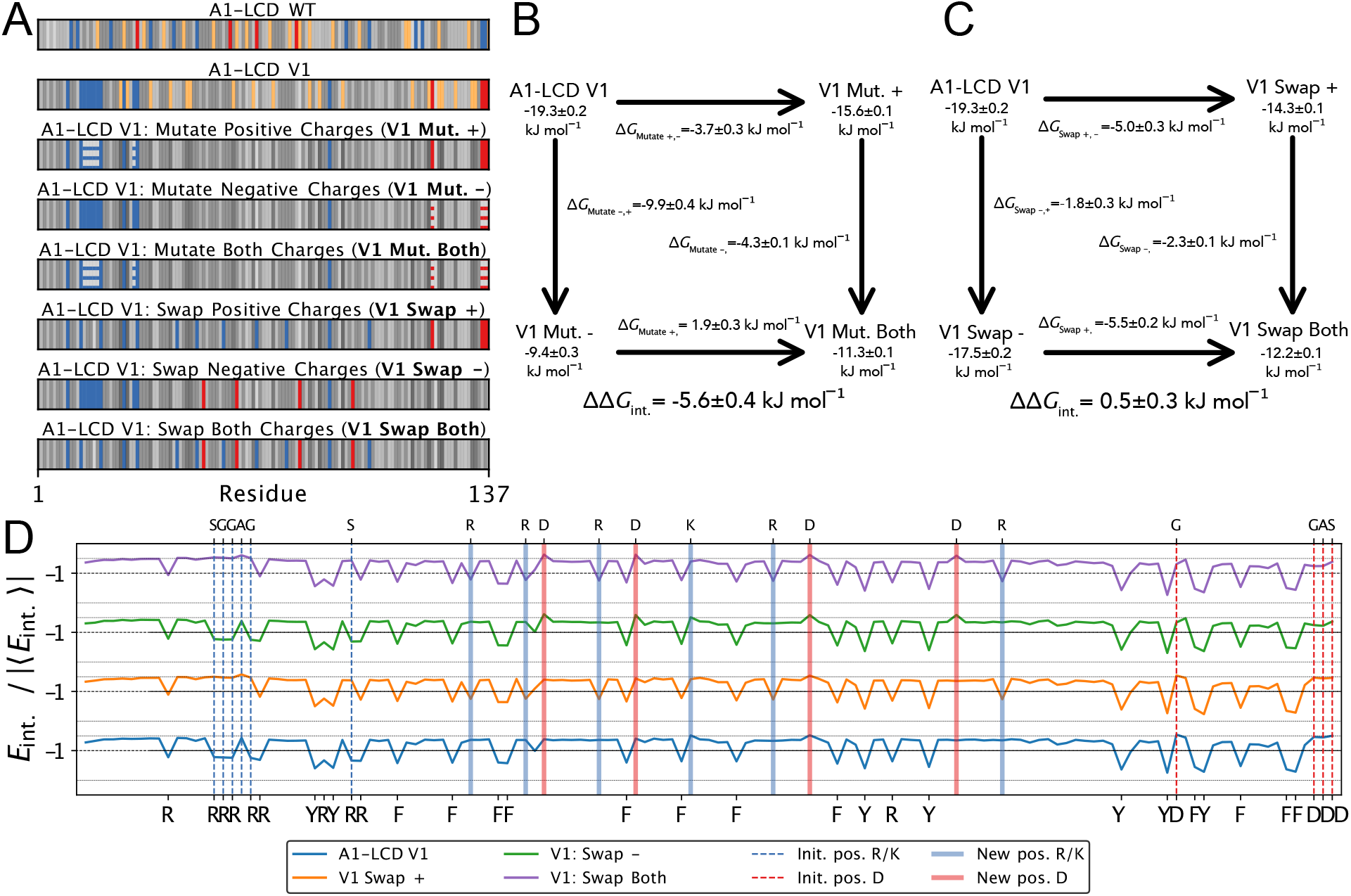
Probing charge interactions with double mutant cycles. (A) Sequence plots the V1 A1-LCD variant (***Pesce et al., 2024***) and charge patterning perturbation variants. The neutral amino acids are coloured in greyscale to represent their hydropathy based on the CALVADOS 2 *λ* parameters. The solid orange lines represent the tyrosine and phenylalanine residue positions, and the dashed orange lines are where any of those are substituted with alanine residues. (B) Double mutant cycle for perturbing charge patterning of the V1 variant with substitutions to alanine. (C) Double mutant cycle for perturbing the charge patterning of the V1 variant, where charged residues near the termini are swapped with non-charged residues internally in the sequence. (D) From the bottom to top, 1D projections of the energy maps for (blue) the V1 variant, V1 where (orange) the positive residues, (green) the negative residues, and (purple) both are swapped back into the sequence. The dashed lines indicate the original positions of the (blue) positive and (red) negative residues, and the traces indicate the new positions. At the bottom, we show the sequence positions of the aromatic and charged residues, and on the top, we show the new positions of the swapped residues. The original maps are normalised by the absolute average interaction energy | ⟨ *E*⟩ |.

We performed slab simulations of V1 and the six variants at 298 K to determine the energetics for the double mutant cycles (Fig. 4B and C). We use this higher temperature (the simulations above were all performed at 278 K), as the increased PS propensity of the V1-variant made the sampling of the dilute phase challenging at 278 K. In the double mutant cycle where we change the net charge of the protein, we observe a greater effect of removing the four negatively charged residues (–9.9 kJ mol^−1^) compared to removing the six positively charged residues (–3.7 kJ mol^−1^) (Fig. 4B). In the case of substituting the positive residues, the net charge becomes more neutral (V1 has a formal net charge of +8 and V1-Mut. + has a formal net charge of +2); this decrease in NCPR thus helps balance out the loss of four arginine sticker residues. In contrast, while changing four aspartates to alanines does not affect the overall stickiness much, the formal net charge increases to +12, thus substantially decreasing PS (***Tesei et al., 2021; Bremer et al., 2022***). The double mutant has a formal net charge of +6 and is thus closer to neutral than V1 (Table S2); this decrease in the NCPR thus helps offset the loss of the sticky arginine residues as well as the potential from long-range charge–charge interactions. This combination of effects means that the effect of substituting five arginines and one lysine with six alanines can be either destabilising (–3.7 ± 0.3 kJ mol^−1^ when introduced into the V1 background) or stabilising (+1.9 ± 0.3 kJ mol^−1^ when introduced into the V1 Mut. - background). Thus, when we calculate the coupling energy, we find that it is negative (–5.6 ± 0.4 kJ mol^−1^). This formally means that the interaction is unfavourable, though this result is the combination of the effects of changing the charge-interactions between the terminal regions and the interaction with the overall net charge of the protein.

To probe the interactions of the charge blocks in the V1 variant while avoiding the complications of changing the overall net charge, we followed a different strategy. As detailed above, we instead changed the location of the positively and negatively charged residues at the N- and C-terminal regions, respectively, by swapping their positions with glycine, serine or alanine residues internally in the sequence. In this way, we conserve the NCPR but disrupt the charge blocks that help stabilise V1 condensates. We again performed direct co-existence simulations to quantify the driving forces for phase separation (Fig. 4C). We find that moving the charged residues from the termini to internal positions decreases phase separation, and that moving the location of six positively charged residues has a greater effect (–5.0 ± 0.3 kJ mol^−1^) than moving four negatively charged residues (–1.8 ± 0.3 kJ mol^−1^). However, the effects of these substitutions are essentially independent of the background so that the resulting coupling energy is close to zero (0.5 ± 0.3 kJ mol^−1^). This, in turn, means that while the sequence separation of positively and negatively charged residues contributes substantially to PS (***Pesce et al., 2024***), the effects of placing the positive residues near the N-terminus and the negative residues near the C-terminus are essentially additive. We note also the variant where we substitute the positive residues, including five sticky arginines (V1 Mut. +), phase separates slightly more strongly than the corresponding swap variant (V1 Swap +), likely because the lower NCPR compensates the loss of the sticky arginine residues in the V1 Mut. + variant. We again analysed the residue-level changes to the interaction energies (Fig. 4D, S5) and S6. For both cycles, we see that in the single mutant steps there is a loss in stabilisation of non-ionic interactions between residues in either N- or C-terminal positions (Figs. S5EF and S6EF), which is not visible in the double mutant step (Figs. S5GH and S6GH). In both the single and double mutant steps, where we perturb the positively charged residues, mainly arginine residues, the effect of the loss of sticky residues is clear. When we swap the residues to internal positions, it is clear that these residues form new interactions in their new positions. The difference map for the substitutions of the positively charged residues also provides insight into the mechanism underlying the increase in PS propensity, coinciding with the change in net charge. While there is a loss of interactions at the positions with substituted arginine residues, sticker residues in other positions are involved in an increased number of interactions. For the swap variants, we find that when we swap the positive residues, the change in interactions is generally similar, going from WT to the single mutant and going from the single mutant, where the negative residues have been swapped already, to the double mutant.

## Conclusions

Here, we set out to explore whether double mutant cycle experiments (***Jencks, 1981; Horovitz, 1996***) could potentially be applied to determine energetic contributions of specific interactions to the PS propensity of an IDR. Such experiments could provide useful information on the nature of the interactions that drive PS and help provide a more detailed mechanistic understanding of previous mutational experiments. To this end, we performed simulated double mutant cycle experiments of A1-LCD variants, attempting to quantify the potential contributions of different types of interactions to the PS propensity and condensate stability. We initially probe the potential of hybrid cation–*π*/*π*–*π* interactions between Arg and Tyr residues that have been suggested to be important for the stability of condensates of PLDs. In our analysis, we find no significant coupling energy (Fig. 2), suggesting these interactions might contribute additively to the PS propensity; this observation is in line with the observed correlation between the number of tyrosine and arginine residues and *c*_sat_ (***Wang et al., 2018***). Next, we probe the *π*–*π*-interactions between aromatic sticker residues, and again observe no measurable couplings when we perturb the aromatic residues at the termini or in the core of the sequence (Fig. 3). Finally, we interrogate charge–charge interactions for the V1 A1-LCD variant with segregated charges and strong electrostatic interactions between the termini. The situation here is more complex because of the effects of changing both the net charge of the protein and the more specific charge–charge interactions. Thus, we observe a negative coupling energy when we substitute the charged residues with neutral alanine residues and thereby change the net charge. In contrast, when we disrupt the charge blocks by moving the charged residues to internal positions, we find essentially additive contributions. Overall, the lack of coupling in the double mutant cycles for the Tyr–Arg interaction, *π*–*π*-interactions and the charge–charge interactions might suggest that many types of interactions in the condensate are additive.

One limitation of our study is that the individual interactions in the CALVADOS model are additive and that the non-ionic (sticky) interactions are described by parameters that are calculated using combination rules using the average ‘stickiness’-parameters (*λ*) and bead sizes (*σ*) of the interacting amino acid residues. Nevertheless, our results with the charge-modifying variants of V1 show a sizeable coupling energy (Fig. 4B), demonstrating that our setup can indeed reveal non-additive effects. Further, we note that it has previously been shown that free energy calculations for double mutant cycles for folding stability using (additive) all-atom force fields can yield sizeable couplings that correlate with experimental measurements (***Werner et al., 2021***).

Another important point is that residue-level coarse-grained models such as CALVADOS can not distinguish between more specific contributions, such as from backbone hydrogen bonds or amide *π*-interactions, and side-chain interactions (***Zeng and Pappu, 2025***). This is a fundamental limitation of coarse-grained models that combine different physical effects into effective interaction parameters. Thus, it is clear that these simulations alone cannot be used to disentangle different physical contributions. More detailed models exist that have multiple interaction sites for each amino acid (***Baul et al., 2019; Souza et al., 2021; Zhang et al., 2024***), and these may aid in separating individual physical contributions.

Looking ahead, we envision that a double mutant cycle approach, as we have presented it here, could be a means of enhancing the interpretation of mutational studies of biomolecular condensates, providing more quantitative insights regarding the interactions stabilising condensates. We also envision that data generated through double mutant cycles could help in deciding which terms and parameters to include when building coarse-grained models, and could provide data for additional benchmarking and parameterisation. For example, one group of coarse-grained models represent the interactions between the 20 amino acids by 210 individual pair terms (***Norgaard et al., 2008; Dignon et al., 2018a; Tesei et al., 2021***), whereas other models represent the same interactions using 20 parameters and combining rules (***Joseph et al., 2021***). While it is clear that a large number of parameters provides additional flexibility in terms of describing individual interactions, it is also more challenging to parametrise. In the context of folded proteins, it has been shown that a full 210-parameter model can be effectively expressed using combinations of 20 amino-acid-specific parameters (***Li et al., 1997; Cieplak et al., 2001***). Similarly, in a top-down parameterisation of a coarse-grained model, it was shown that going from 20 to 210 parameters only led to a small improvement in agreement with experiments (***Dannenhoffer-Lafage and Best, 2021***). Nevertheless, other models have shown that adding specific pair terms can help capture specific types of interactions (***Martin et al., 2020; Das et al., 2020; Joseph et al., 2021; Bremer et al., 2022***), and we hope that double mutant cycle studies can help provide data for testing and parametrising such more detailed models.

Finally, while we have here explored the effects of interactions on condensate stability in the form of a partitioning free energy of protein chains, it would also be possible to probe other properties of biomolecular condensates, such as material properties or kinetics of exchange. Double mutant cycles could potentially also be used to build and test hypotheses about interactions between different types of proteins and nucleic acids, or for interactions between IDRs and folded domains.

## Methods

### Molecular Dynamics simulations

We performed all MD simulations with the CALVADOS 2 force field (***Tesei and Lindorff-Larsen, 2023; von Bülow et al., 2025***) in the NVT ensemble using a Langevin integrator with a 10 fs time step and a 0.01 ps^−^1 friction coefficient from the openmm package (v 7.5) (***Eastman et al., 2017***). We model the bonded interactions with a harmonic potential,

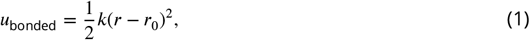

where force constant *k* = 8030 kJ mol^−1^nm^−2^ and an equilibrium bond length *r*_0_ = 0.38 nm. We model the non-ionic non-bonded interactions with a 12-6 Lennard-Jones potential,

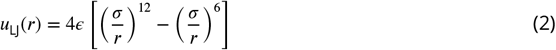

which we use in a truncated and shifted Ashbaugh-Hatch potential (***Ashbaugh and Wood, 1997; Ashbaugh and Hatch, 2008***),

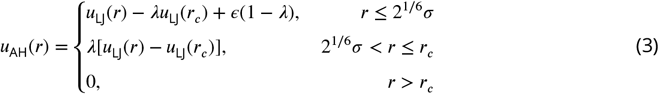

where *ε* = 0.8368 kJ mol^−1^, the truncation cut-off *r* = 2 nm, *λ* is the arithmetic average of hydropathy values of two interacting residues, and *σ* is the arithmetic average of the Van der Waals diameters of the interacting residues. The amino-acid specific *σ*-values have been determined by ***Kim and Hummer*** (***2008***). We model the salt-screened ionic interactions with a truncated and shifted Debye-Hückel potential with a cut-off *r*_*c*_ = 4 nm,

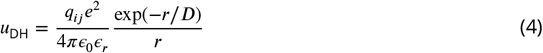

where *q*_*ij*_ are the multiplied charge numbers of the interaction residues, *e* is the elementary charge, *ϵ*_0_ is the vacuum permittivity, and 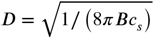 is the Debye-length of an electrolyte solution of ionic strength, *c*_*s*_, and Bjerrum length, *B*. We model the dielectric constant of the implicit aqueous solution, *ϵ*_*r*_, in a temperature-dependent manner with an empirical relationship (***Akerlof and Oshry, 1950***),

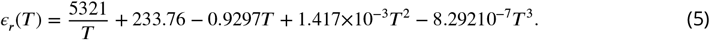

### Setup and analysis of slab simulations

To model phase separation, we performed direct-coexistence simulations in an elongated rectangular simulation box with dimensions *l*_*s*_ *>> l*_*x*_ = *l*_*y*_. In this approach, we use the periodic boundaries to form a slab geometry (***Blas et al., 2008; Silmore et al., 2017; Dignon et al., 2018a; Tesei et al., 2021***). For the dimensions of the box and the number of protein chains, we chose to use the same values as ***Tesei et al***. (***2021***) (*N*_mol_, *l*_*xy*_ = 15 nm and *l*_*s*_ = 150 nm). We set up the starting topology of systems by placing the protein chains in the centre of the simulation box as non-overlapping Archimedean spirals, where the beads of one chain are spaced by the equilibrium bond length. All simulations have been performed at 278 K, except for the A1-LCD ‘V1’ variant-based systems, which were performed at 298 K. The ionic strength (*c*_*s*_) is set to 150 mM in all simulations. We determined boundary *z*-values for the phases from the equilibrium concentration profile along the *z*-axis to calculate the concentrations in the dense- and dilute-phases. As described in previous work, this profile is determined after first centring the protein-dense slab in the box (***Jung and Yethiraj, 2018; Tesei and Lindorff-Larsen, 2023; von Bülow et al., 2025***). We report the *z*-axis boundaries used in this study in a DataFrame deposited in the GitHub repository supporting this work.

To quantify a protein’s propensity to phase separate, we calculate Δ*G*_trans_,

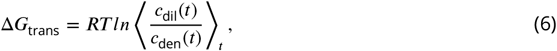

where *R* is the gas constant and *c*_dil_(*t*) and *c*_den_(*t*) are the time series of the dilute- and dense-phase concentrations, respectively.

### Calculation of Double Mutant Cycles

For calculations of double mutant cycles, we use the implementation as outlined by ***Horovitz*** with Δ*G*_trans_ as our measure of phase separation process, which we can determine from our simulations.

We calculate the difference in free energy for the single mutant steps, Δ*G*_P–XY→P–xY_ and Δ*G*_P–XY→P–Xy_ and adding the subsequent double mutant steps, Δ*G*_P–Xy→P–xy_ and Δ*G*_P–xY→P–xy_ as follows from the simulations,

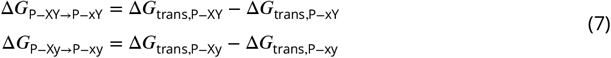

where X and Y represent the initial residues, x and y the mutated mutated residues and Δ*G*_trans_ is calculated from the simulations as described (Eq. 6).

To quantify any potential coupling between the perturbed amino acid residues, we calculate the difference between the energy change for the single and the double mutant steps for the separate residues,

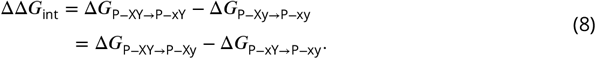

## Data and Code Availability

All the data and code used for this work are available via https://github.com/KULL-Centre/_2026_rauh_dmc. The CALVADOS software package (***von Bülow et al., 2025***) is available at https://github.com/KULL-Centre/CALVADOS.

## Acknowledgements

We thank Sören von Bülow and other members of SBiNLab for helpful discussions. We acknowledge access to computational resources from the ROBUST Resource for Biomolecular Simulations (supported by the Novo Nordisk Foundation grant no. NF18OC0032608) and the Danish National Supercomputer for Life Sciences (Computerome). This work is a contribution from the PRISM (Protein Interactions and Stability in Medicine and Genomics) centre funded by the Novo Nordisk Foundation (to K.L.-L.; NNF18OC0033950).

## Supplemental Information for

**Figure S1.**
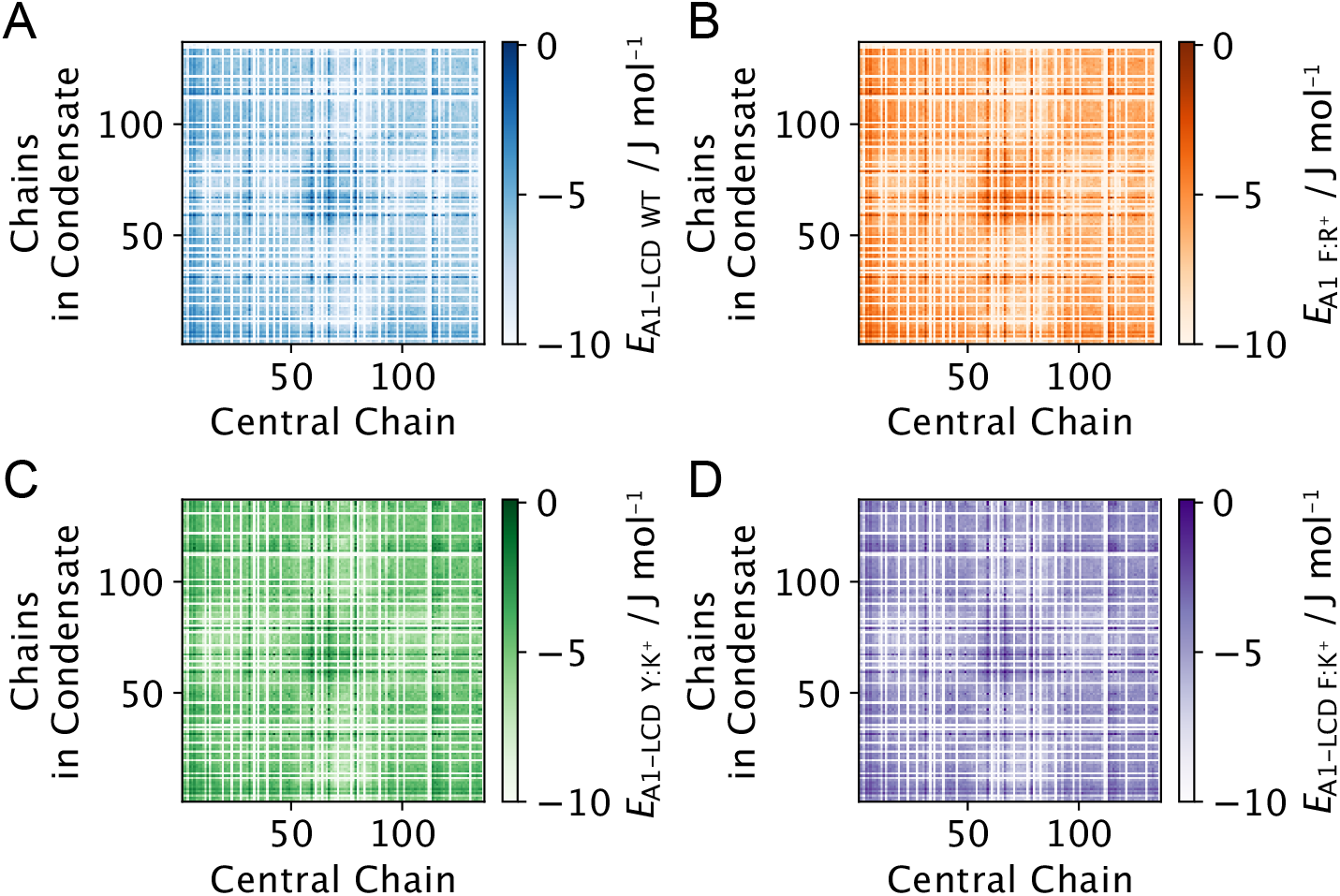
Interaction maps for the YR-FK cycle. Residue–residue interaction maps for non-ionic interactions for (A) A1-LCD WT, (B) A1-LCD Y → F, (C) A1-LCD R → K, and (D) the A1-LCD double mutant with Y → F and R → K

**Figure S2.**
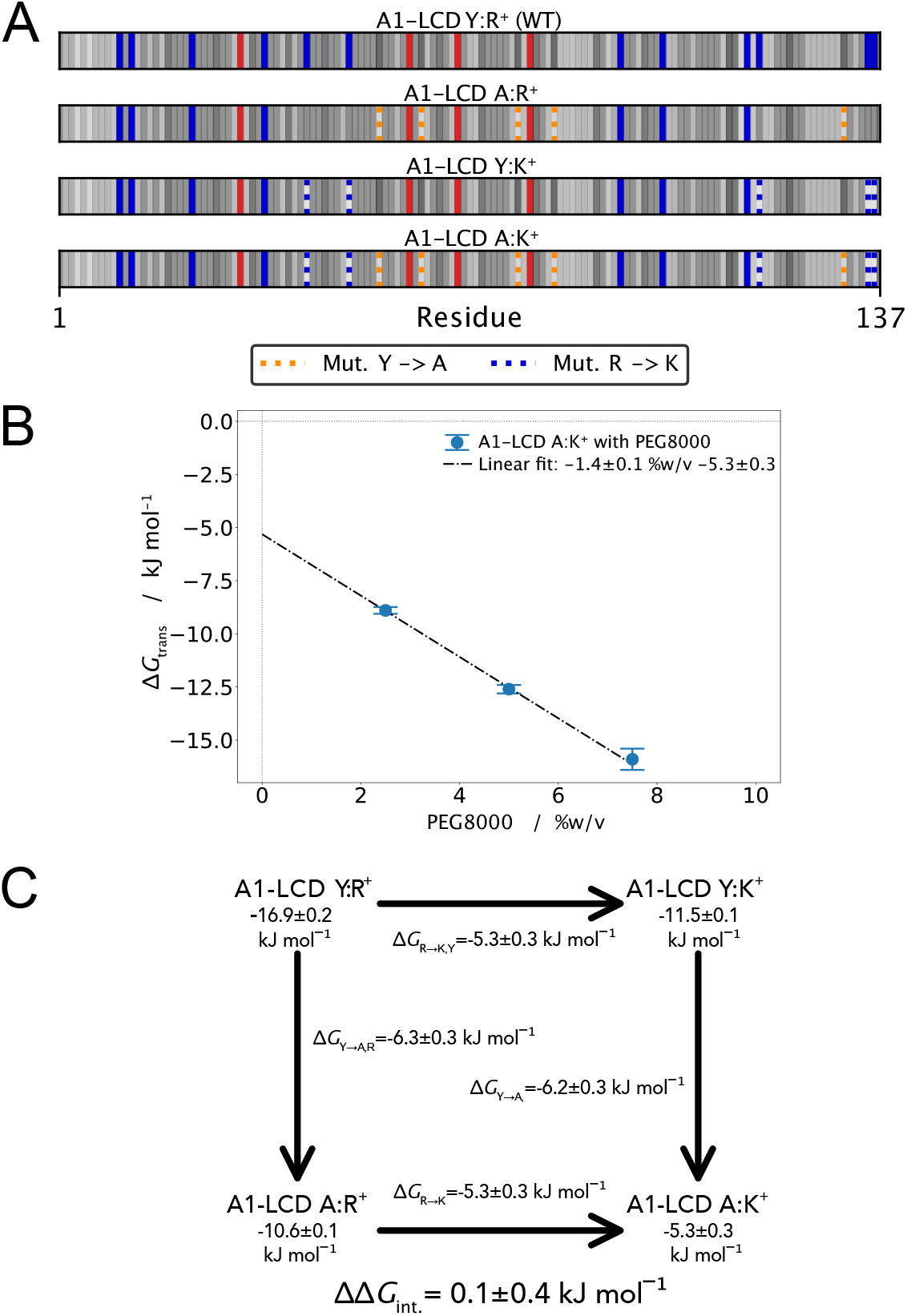
Alternative cycle for tyrosine–arginine interactions. (A) Sequence plots of A1-LCD WT and variants. The blue and red lines represent positively and negatively charged amino acid residues. The neutral amino acids are coloured in greyscale to represent their hydropathy based on the CALVADOS 2 *λ* parameters. The dashed lines represent the mutations of (orange) tyrosine to alanine residues and (blue) arginine to lysine residues. (B) Simulated PEG titration curve for A1-LCD A:K^+^ (C) Double mutant cycle for perturbing Arg–Tyr interactions with the mutations R→K and Y→A.

**Figure S3.**
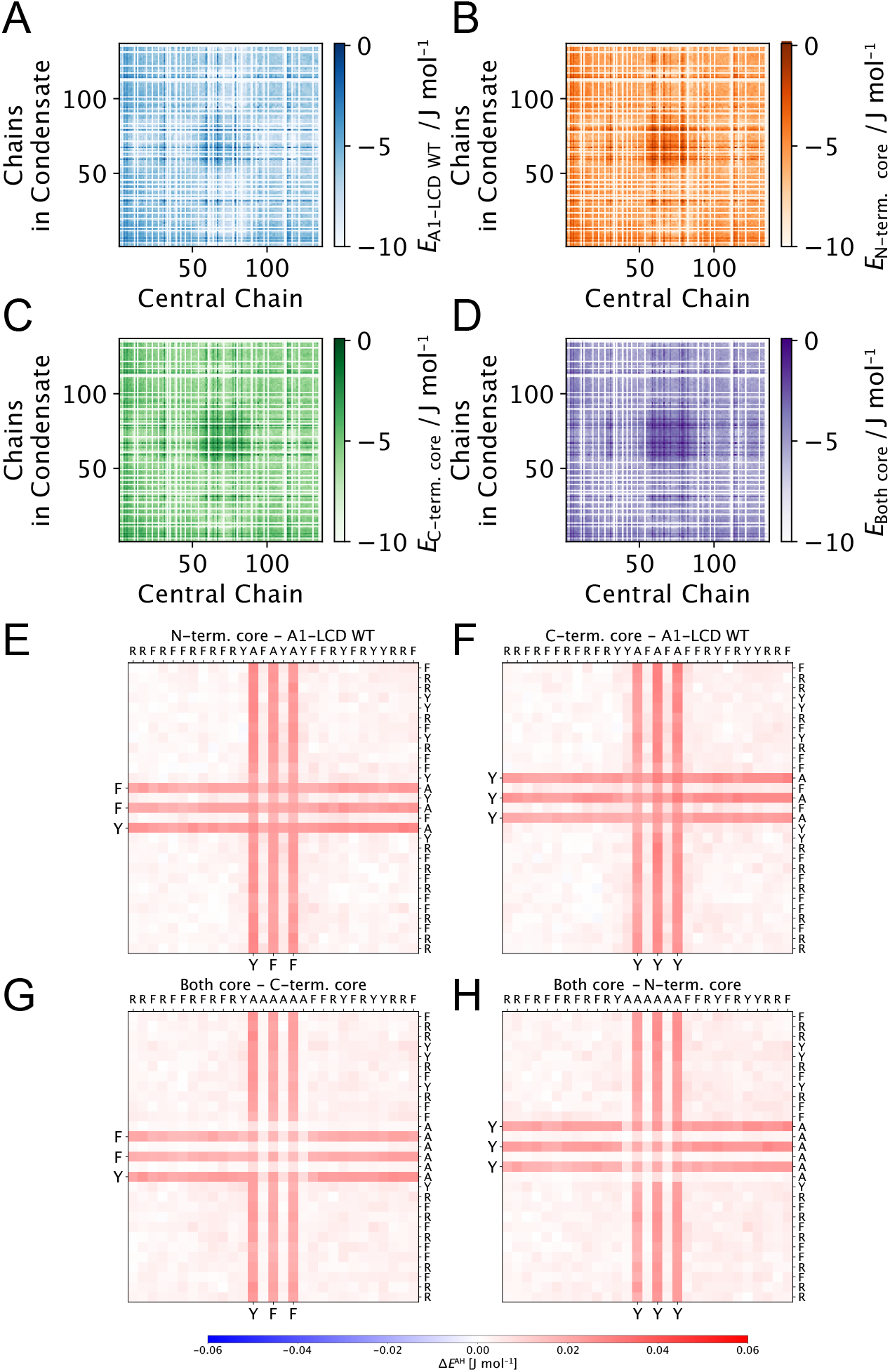
Effect of perturbing sticker residues in the central core region. Residue–residue interaction maps for non-ionic interactions for (A) A1-LCD WT, (B & C) A1-LCD, where two different sets of aromatic residues in the core are mutated (orange & green), and (D) A1-LCD, where all core aromatics are mutated (purple). Sticker residue interaction difference map for (E) mutating the N-term. core residues, (F) mutating the N-term. core residues, (G) mutating the N-term. core residues after the C-term. core residues, and (H) mutating the C-term. core after the N-term. core residues.

**Figure S4.**
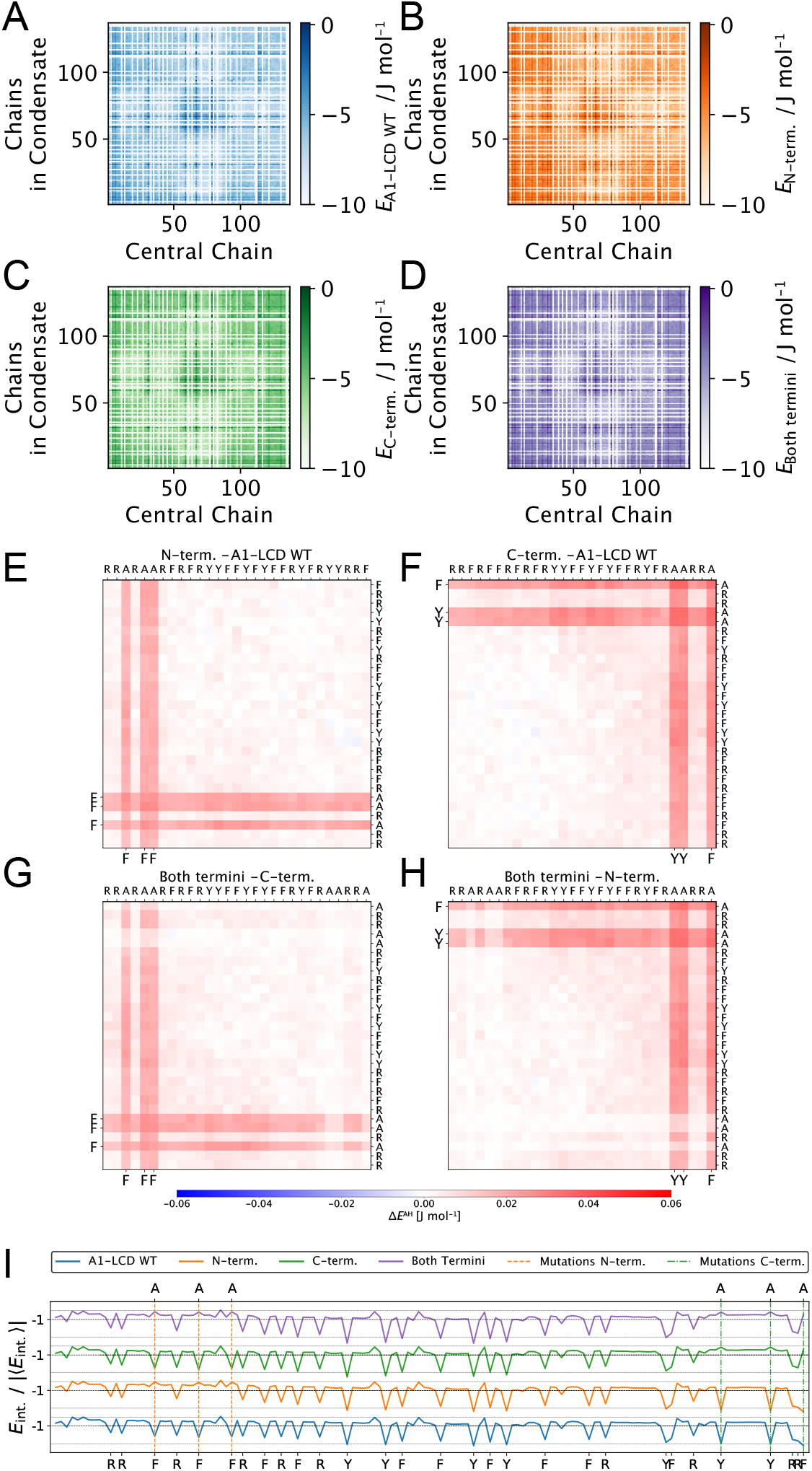
Effect of perturbing sticker residues near the termini. Residue–residue interaction maps for non-ionic interactions for (A) A1-LCD WT, A1-LCD, where aromatic residues are either (B) at the N-terminal region or (C) at the C-terminal region are mutated (orange & green), and (D) A1-LCD, where both the N-terminal and C-terminal aromatics are mutated (purple). Sticker residue interaction difference map for mutating the N-terminal residues, (F) mutating the C-terminal residues, (G) mutating the N-terminal residues in the context of mutated C-terminal residues, and (H) mutating the C-terminal residues in the context of mutated N-terminal residues. (I) From the bottom to top, 1D projections of the energy maps for (blue) A1-LCD WT, (orange & green) A1-LCD where two different sets of aromatic residues at the termini are mutated, and (purple) A1-LCD where all termini aromatics are mutated. The maps are normalised by the absolute average interaction energy |⟨*E*⟩|

**Figure S5.**
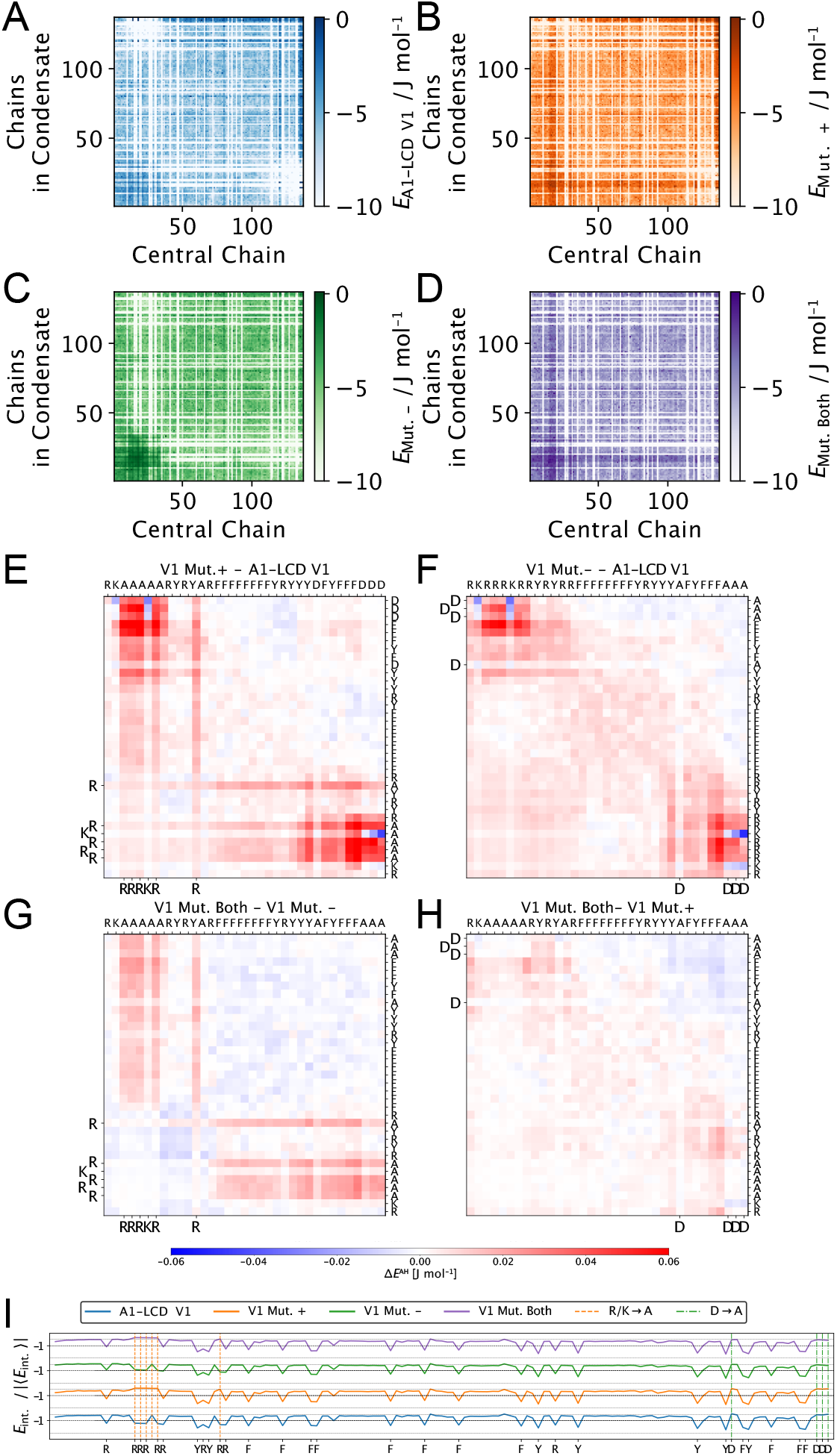
Effect of perturbing charge interactions in V1 while also changing the net charge. Residue–residue interaction maps for non-ionic interactions for (A) A1-LCD V1 variants, (B) A1-LCD V1 mutating positively charged residues to alanine residues, (C) A1-LCD V1 mutating negatively charged residues to alanine residues, and (D) A1-LCD V1 mutating both positively and negatively charged residues to alanine residues. Sticker residue interaction difference maps for mutating (E) positively charged residues, negatively charged residues, (G) positively charged residues in a context where the negative residues have been removed, and (H) negatively charged residues in a context where the positively charged residues have been removed. (I) From the bottom to top, 1D projections of the energy maps for (blue)A1-LCD swap variant V1, (orange) V1 where the positive residues and (green) the negative residues and (purple) both are mutated to alanine residues. The dashed lines indicate the original positions of the (blue) positive and (red) negative residues. At the bottom, we show the sequence positions of the aromatic and charged residues. The maps are normalised by the absolute average interaction energy |⟨*E*⟩|.

**Figure S6.**
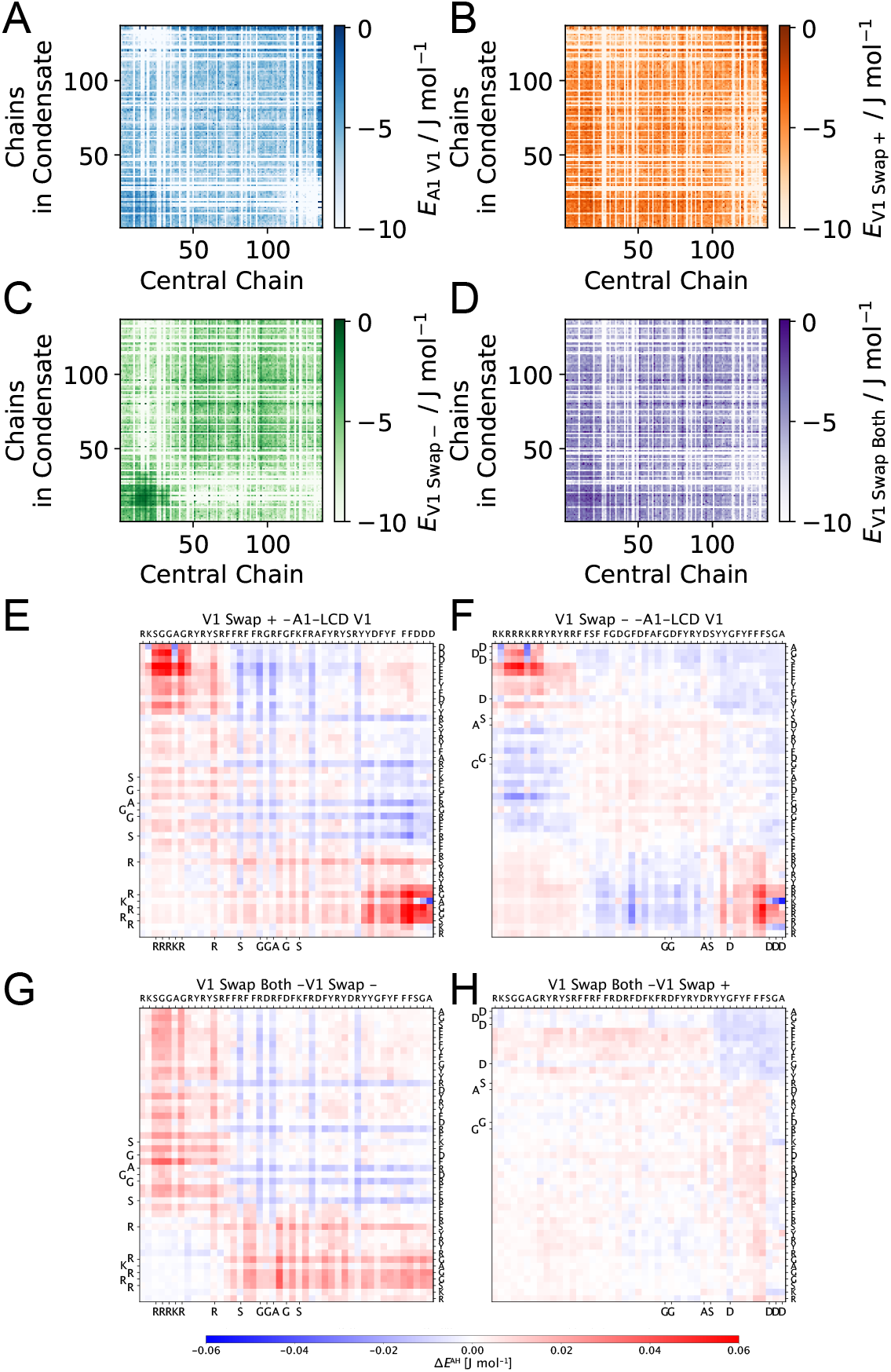
Effect of perturbing charge interactions in V1 while maintaining the net charge. Residue–residue interaction maps for non-ionic interactions for (A) A1-LCD V1, (B) A1-LCD V1 swapping positively charged residues, (C) A1-LCD V1 swapping negatively charged residues, and (D) A1-LCD V1 swapping both positively and negatively charged residues. Sticker residue interaction difference map for swapping (E) positively charged residues, (F) negatively charged residues, (G) positively charged residues where negatively charged residues have been swapped, (H) negatively charged residues where positively charged residues have been swapped.

**Table S1.**
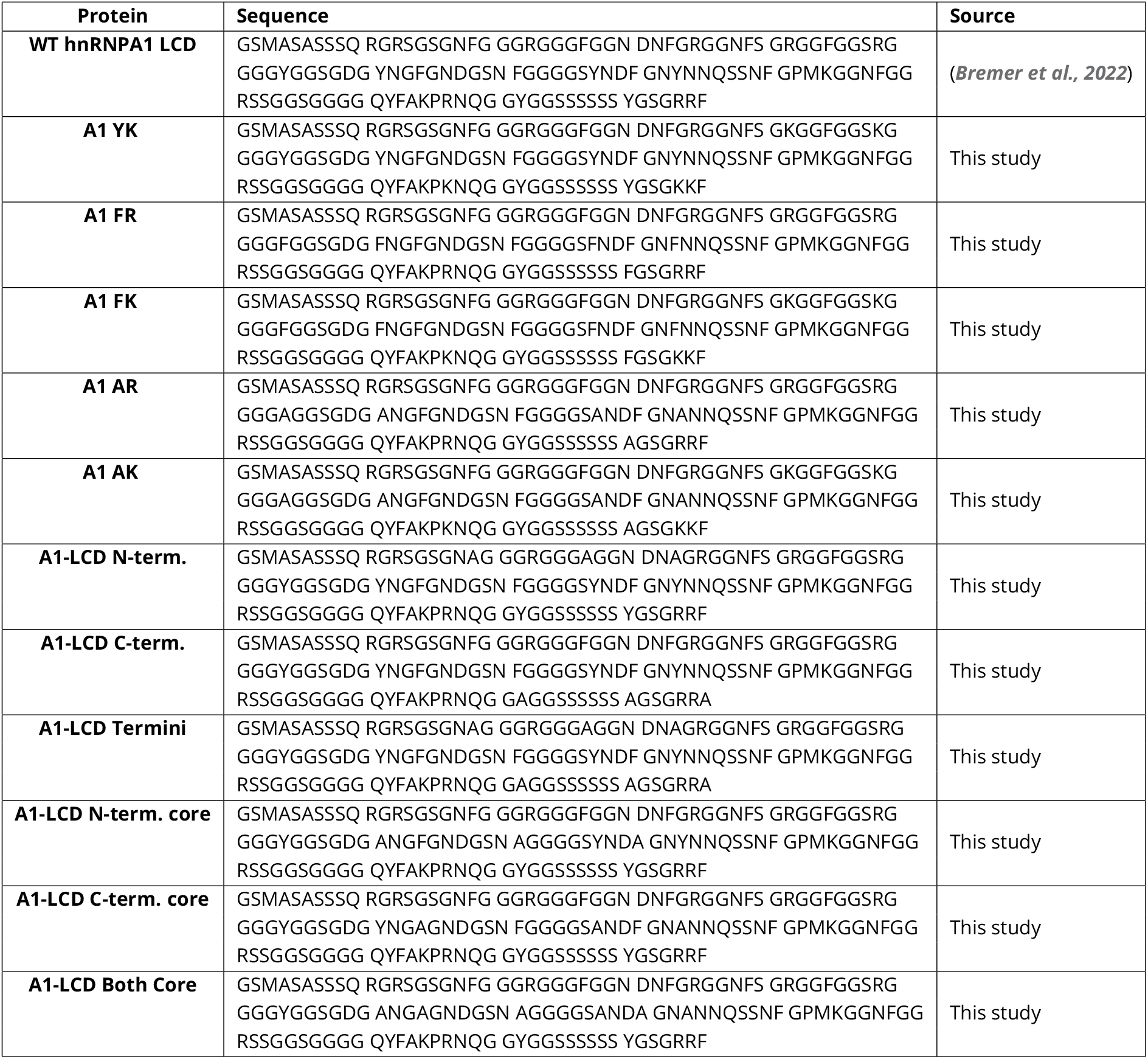
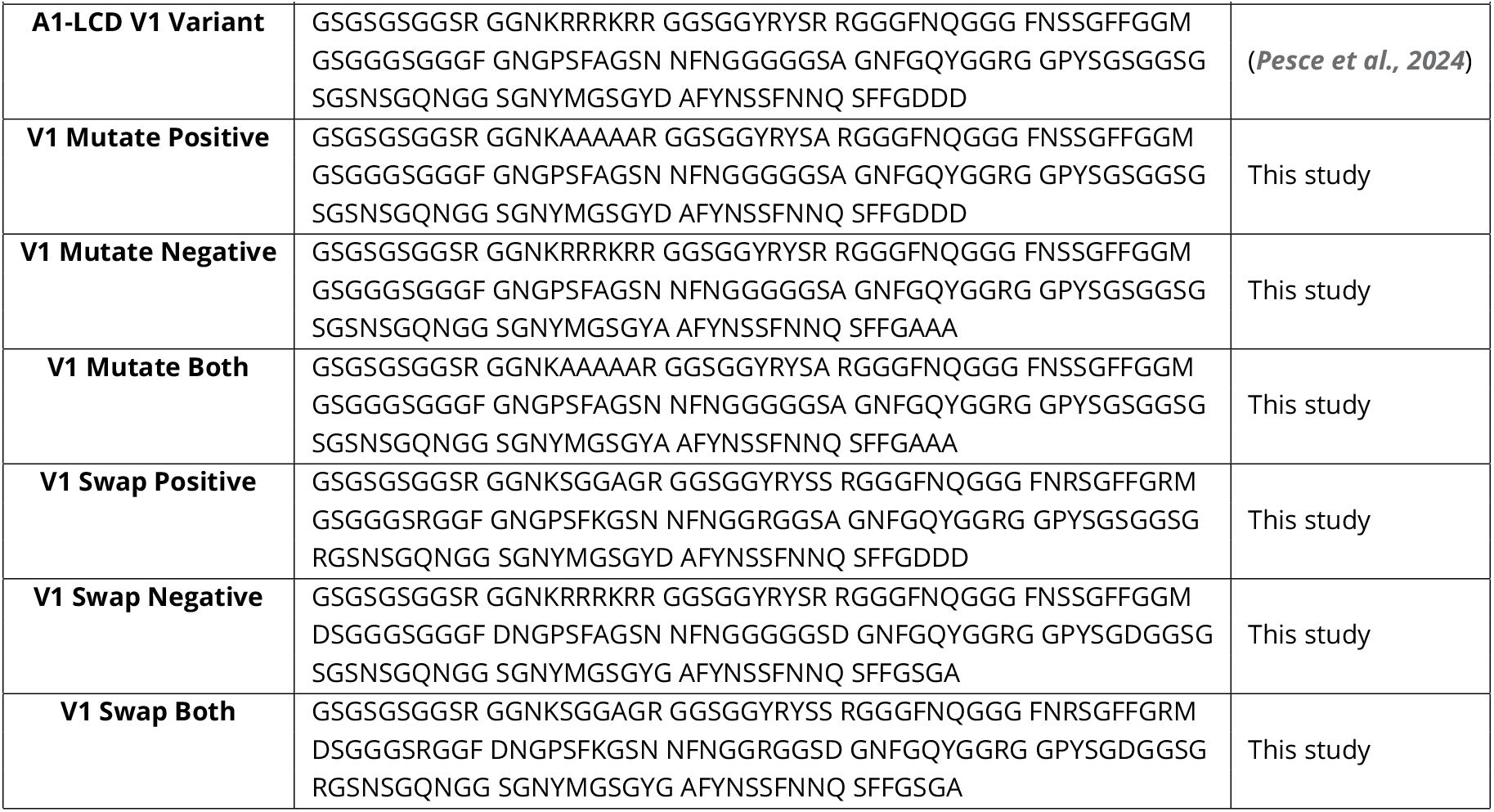
Sequences of proteins simulated in this study.

**Table S2.** Sequences descriptors calculated for the sequences of proteins and variants simulated in this study. *f*_K_, *f*_R_, *f*_E_, and *f*_D_ are the fractions of lysine, arginine, glutamate and aspartate residues, respectively.

| Protein | $\langle\lambda\rangle$ | SHD | SCD | $\kappa$ | FCR | NCPR | $f_K$ | $f_R$ | $f_E$ | $f_D$ | $f_{aro}$ |
| --- | --- | --- | --- | --- | --- | --- | --- | --- | --- | --- | --- |
| WT A1-LCD | 0.611 | 5.506 | 1.337 | 0.183 | 0.131 | 0.058 | 0.015 | 0.073 | 0.0 | 0.029 | 0.139 |
| A1-LCD Y:K<sup>+</sup> | 0.591 | 5.34 | 1.337 | 0.183 | 0.131 | 0.058 | 0.051 | 0.036 | 0.0 | 0.029 | 0.139 |
| A1-LCD F:R<sup>+</sup> | 0.607 | 5.469 | 1.337 | 0.183 | 0.131 | 0.058 | 0.015 | 0.073 | 0.0 | 0.029 | 0.139 |
| A1-LCD F:K<sup>+</sup> | 0.587 | 5.302 | 1.337 | 0.183 | 0.131 | 0.058 | 0.051 | 0.036 | 0.0 | 0.029 | 0.139 |
| A1-LCD A:R<sup>+</sup> | 0.586 | 5.269 | 1.337 | 0.183 | 0.131 | 0.058 | 0.015 | 0.073 | 0.0 | 0.029 | 0.102 |
| A1-LCD A:K<sup>+</sup> | 0.565 | 5.102 | 1.337 | 0.183 | 0.131 | 0.058 | 0.051 | 0.036 | 0.0 | 0.029 | 0.102 |
| A1-LCD N-term. | 0.598 | 5.388 | 1.337 | 0.183 | 0.131 | 0.058 | 0.015 | 0.073 | 0.0 | 0.029 | 0.117 |
| A1-LCD C-term. | 0.597 | 5.397 | 1.337 | 0.183 | 0.131 | 0.058 | 0.015 | 0.073 | 0.0 | 0.029 | 0.117 |
| A1-LCD Termini | 0.584 | 5.279 | 1.337 | 0.183 | 0.131 | 0.058 | 0.015 | 0.073 | 0.0 | 0.029 | 0.095 |
| A1-LCD N-term. core | 0.597 | 5.374 | 1.337 | 0.183 | 0.131 | 0.058 | 0.015 | 0.073 | 0.0 | 0.029 | 0.117 |
| A1-LCD C-term. core | 0.597 | 5.366 | 1.337 | 0.183 | 0.131 | 0.058 | 0.015 | 0.073 | 0.0 | 0.029 | 0.117 |
| A1-LCD Both Core | 0.583 | 5.234 | 1.337 | 0.183 | 0.131 | 0.058 | 0.015 | 0.073 | 0.0 | 0.029 | 0.095 |
| A1-LCD V1 | 0.611 | 5.539 | -2.651 | 0.536 | 0.131 | 0.058 | 0.015 | 0.073 | 0.0 | 0.029 | 0.139 |
| V1 Mutate Positive | 0.595 | 5.398 | -1.674 | 0.257 | 0.088 | 0.015 | 0.007 | 0.036 | 0.0 | 0.029 | 0.139 |
| V1 Mutate Negative | 0.618 | 5.586 | 1.113 | 0.575 | 0.102 | 0.088 | 0.015 | 0.073 | 0.0 | 0.0 | 0.139 |
| V1 Mutate Both | 0.602 | 5.445 | 0.232 | 0.215 | 0.058 | 0.044 | 0.007 | 0.036 | 0.0 | 0.0 | 0.139 |
| V1 Swap Positive | 0.611 | 5.541 | -0.922 | 0.18 | 0.131 | 0.058 | 0.015 | 0.073 | 0.0 | 0.029 | 0.139 |
| V1 Swap Negative | 0.611 | 5.506 | -1.09 | 0.504 | 0.131 | 0.058 | 0.015 | 0.073 | 0.0 | 0.029 | 0.139 |
| V1 Swap Both | 0.611 | 5.508 | 0.681 | 0.155 | 0.131 | 0.058 | 0.015 | 0.073 | 0.0 | 0.029 | 0.139 |

